# Working out Classic Examples of Metabolic Networks Structural Analysis with Wolfram Language

**DOI:** 10.1101/672881

**Authors:** Kuo Kan Liang

## Abstract

Metabolic reaction networks usually include a large number of enzymatic reaction steps. Nowadays, people desire to work on larger and larger metabolic networks, in the research fields of for example systems or synthetic biology. With the growing sizes of the metabolic networks considered, it is more and more tedious and cumbersome for a human to perform routine processing and analysis of the networks. Computer programs that automate metabolic analysis will be beneficial. In this work, the use of Wolfram Language to automate metabolic analysis is demonstrated. Although there are many different analyses to perform in a metabolic network analysis project, the analysis of metabolic network structure may be the most common first step and one of the most troublesome tasks. Therefore, instead of demonstrating a lot of different analyses, the structures of a few important examples from one chapter in the classic textbook by Professor Gregory Stephanopoulos are analyzed with a program written in Wolfram Language and the models are book-kept in JSON format.

## I. INTRODUCTION

Metabolic engineering is generally defined as the modification of metabolic activities in cells for specific functions, mainly metabolites production, through molecular biological techniques[1]. To this end, theoretical tools for studying the behavior of metabolic pathways are essential. Although the metabolic pathways can be categorized into groups according to their functions, and be investigated and explained separately in literature or textbooks, in living cells metabolic pathways are still coupled with each other. Even if two pathways did not share any enzyme or substrate, they may still be coupled through energetic molecules like ATP, and are only approximately separable from one another under nearly perfectly homeostatic situation. Therefore, to be more useful, theoretical analysis should be able to handle larger and larger reaction networks, which by definition involve many metabolic pathways and thus very many enzymatic reactions.

A reaction pathway involving say one hundred enzymatic reactions may not be much more complicated than a pathway involving say ten enzymatic reactions, if there is only very few “branch point” in the pathway. Or in other words, the number of branch points in a reaction network is the most important factor that determines how complicate that network is. A non-branched path between two branch points can always be treated in simple ways when its detail kinetics is not relevant. Therefore, analyzing the structure of a reaction network so that its branch points can be found out and non-branched paths be grouped is a very important first step that simplifies the network so that analyses of large networks become formidable tasks.

In their classic textbook[1], Professor G. N. Stephanopoulos and coauthors spent one chapter (Chapter 12) to explain how to analyze the structure of metabolic networks. In this work, Wolfram Language program that runs on Mathematica[2] software is used to reproduce the analyses mostly according to the procedures explained in the book. Since the explanations in the book are mostly descriptive, they have to be converted into mathematical or logical operations in the program, for automation. In the following, the details of the conversions from theory to program are explained.

In Section II, the three model networks are shown, together with the JSON files used to describe them to the Mathematica program. In Section III, the way the Mathematica program parse the JSON file, the way the program setup important variables for calculation, and the way these information are converted into the steady-state internal metabolites stoichiometric coefficient matrix (SIMS) and the kernel of this matrix are explained in detail. In Section IV, starting with the kernel of SIMS, the link metabolites of the network is found out step by step, with detailed explanation.

## II. MODEL SYSTEMS AND THEIR REPRESENTATIONS

Three metabolic networks are taken as model systems. The first two networks are abstract models with only six metabolites involved. Since the abstract metabolites are named from A to F, sequentially, these two models are called the first and second ATOF models. However, in order to focus on the topology of the problem, the stoichiometric coefficients were simplified, compared with the original model. The third network, which is realistic, composes of the essential parts of the aromatic amino acid synthesis pathway (AAASP).

In order to input the information and parameters of the networks to the program efficiently and systematically, these systems are described in JSON format.

### A. First ATOF model

The network is depicted in the diagram below:

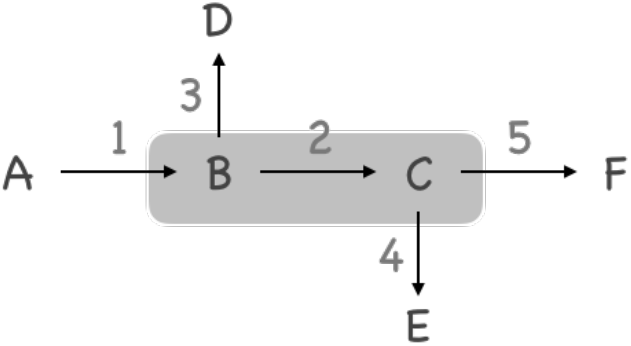

Six abstract metabolites A to F are involved. The arrows represent fluxes among metabolites, and their indices are shown next to them. The order of these fluxes will be respected in the following discussions so the indices are absolute, just like the names of the metabolites. Moreover, since they represent ‘fluxes’ instead of just ‘reactions’, they are, by default, bidirectional (reversible). But in the diagram these arrows are only unidirectional because they represent the presumed direction of those fluxes. Metabolites A, D, E, and F are considered as *terminal metabolites* or *end metabolites* because in further analyses their influxes or effluxes have to be considered. In later text, an influx or an efflux will be called an *exchange flux* for brevity[3]. Metabolites B and C are considered as *internal metabolites*. There is no influxes of them that can be monitored. Consequently, flux number 2 also cannot be directly observed while the pathway is active. Metabolites B and C, together with flux number 2, are enclosed in a light-gray box to show that they are the internal part of the system.

Given the (presumed) directions of the fluxes, the reactions comprising this network can be listed. In this particular simple and abstract model, however, it is too trivial to write them down. In the following, only the JSON file used to describe the network to the Mathematica program is shown.

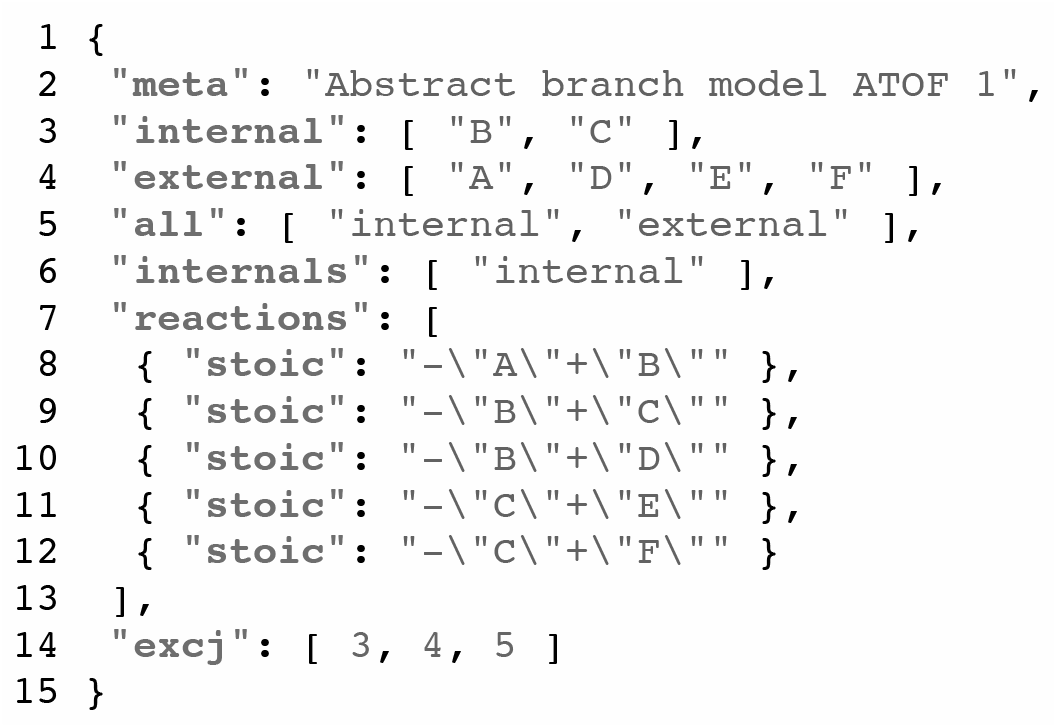

Notice that all of the metabolites (chemical species) are written in strings. There should be a unique name for each metabolite so that later, when we are able to merge two or more networks into one big network, metabolites with the same name should mean the same species across the sub-networks. The *stoichiometric equations* of the fluxes are given. The metabolites are divided into `external` and `internal` groups. There is another group with the key `all`, and its content indicates that the groups `internal` and `external` comprise all of the metabolites in the system. For analyzing the structure of the network, only the internal metabolites are important. However, for this small system, all of the metabolites are included in the `all` group to show how these fields work. At the end of the JSON file there is a record with key `excj`, meaning exchange fluxes. In here three of the four presumed external fluxes are included. Flux number 1 is not included. With hindsight, flux 1 is excluded because it will be divided into flux 2 and flux 3 (therefore B is a branch point). In contrast, flux 3, 4, and 5 are all very simple. In their presumed direction, they will not be further divided into branches. While constructing the kernel (null space) of the steady-state internal metabolites stoichiometric coefficients matrix, this information is crucial. It shall be important to choose proper external fluxes for the analysis to really work. There are other clues with which one can choose the proper external fluxes correctly, but they involve more details of the theory. Therefore they will be explained later.

### B. Second ATOF model

The network is depicted in the diagram below:

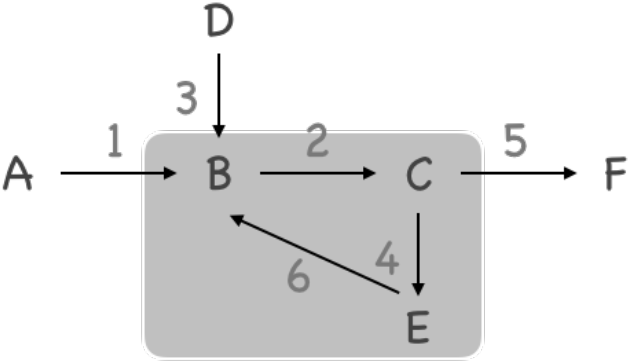

Notice that this network is roughly the same as ATOF model 1, but now the (presumed) direction of flux number 3 is reversed. Besides, there is a flux linking metabolites E and B. this extra flux makes E an internal metabolite. What becomes troublesome in this network is that, in this case, special care has to be taken to find the proper exchange fluxes. To save space, the JSON file describing this model is not shown.

### C. Aromatic amino acid synthesis pathway

This case is based on the model of aromatic amino acid systhesis pathway in yeast published by Galazzo and Bailey[4]. Glucose is used as the only carbon source. Besides the three aromatic amino acids produced, accompanying storage compounds in the form of polysaccharides and glycerol are also included. The free energy flux is considered in the for of ATP to ADP conversion. The diagram of the complete pathway is shown in the following figure.

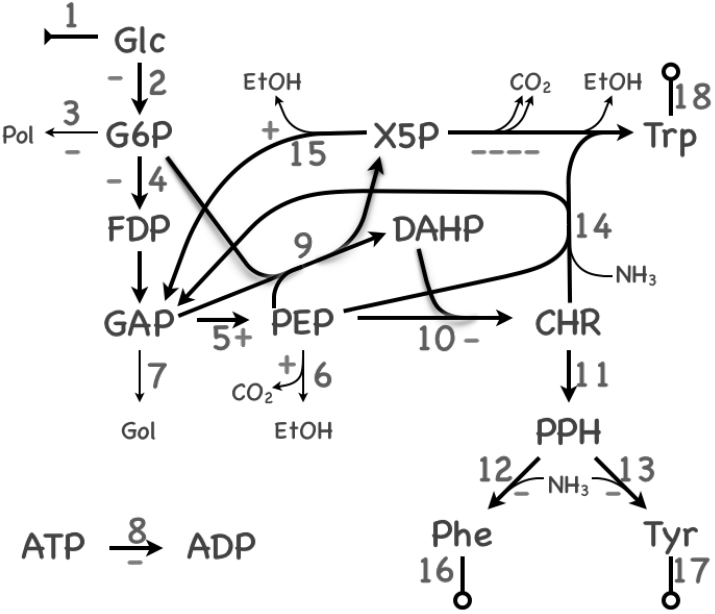

This diagram is much complicated, but the only major difference from the diagrams of simpler and abstract models is that for a flux that produces ATP in its presumed direction, there is a plus sign next to it. If it consumed ATP in its presumed direction, a minus sign is placed next to it. The JSON file for this network model is also shown.

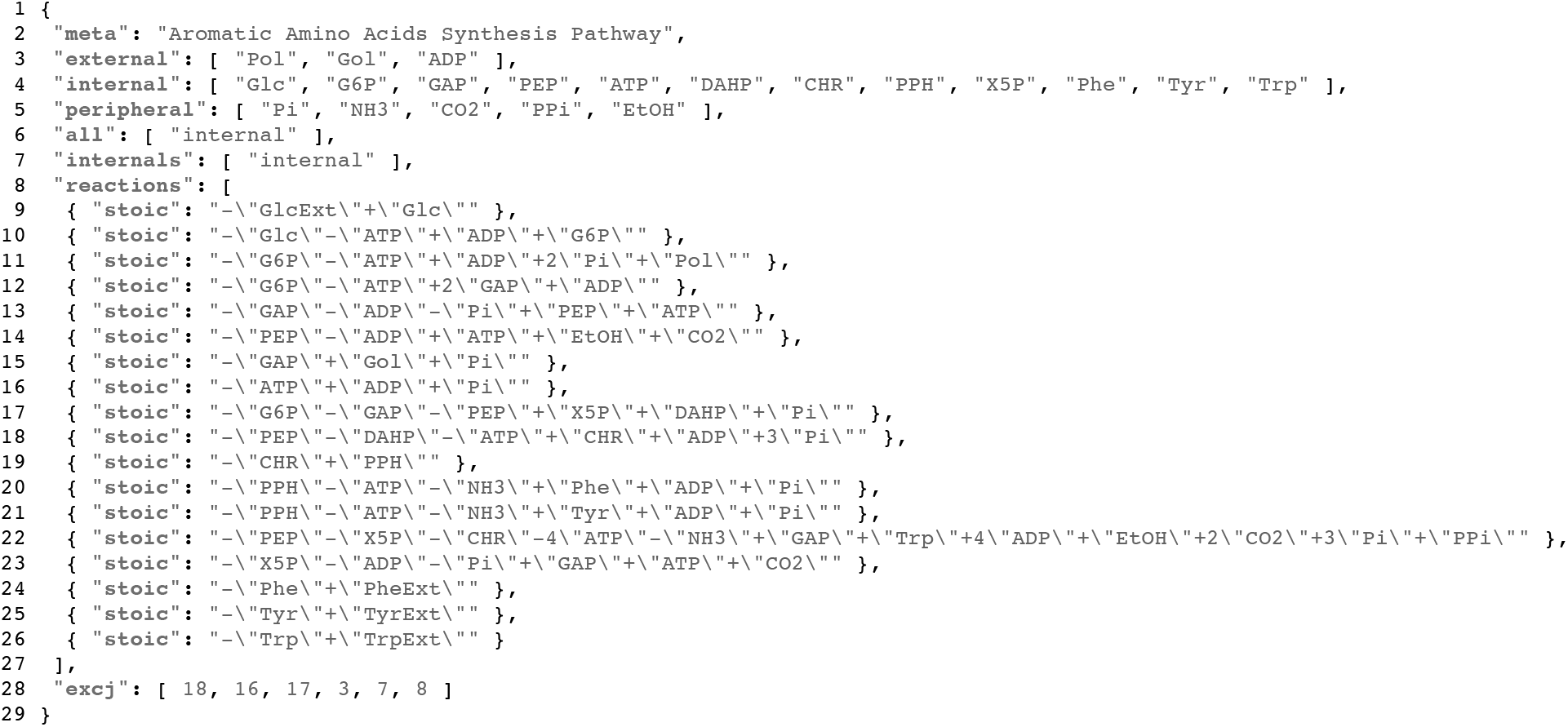

There are totally eighteen fluxes involved. The fluxes are numbered in consistence with the book of Stephanopoulos. However, there are two simplifications. First, although in the diagram flux number 4 is shown to convert G6P into FDP, but since FDP is then converted into two GAP and Stephanopoulos did not consider the flux from FDP to GAP, either, here in the JSON file reaction four the FDP is replaced by two GAP. Second, for maintaining the equilibrium between ATP, ADP, and AMP, in the original model, the reaction linking these three metabolites is also included. However, since it is not related to the analysis of network structure, this flux is ignored in the present work.

## III. INDEPENDENT PATHWAY ANALYSIS

Wolfram Language (or Mathematica. Abbreviated as WL in the following text.) has very good compatibility with JSON notation format. It can import JSON file directly into a nested list. Conversion to list is easy because key-value pairs in JSON can be converted into a rule in WL, and ordered list in JSON can be converted into general list in WL. However, in newer editions of WL, there is a new type of structure called **`Association`** which closely resembles list but more convenient for implementing functionality like dictionary in other languages. Although in the present work most of the functions can also be implemented with list, in other applications it still may be better to use association instead. Therefore we still convert the nested list into nested association. As the first example the JSON file describing ATOF 1 model is imported (suppose that this file is in the working directory of WL):

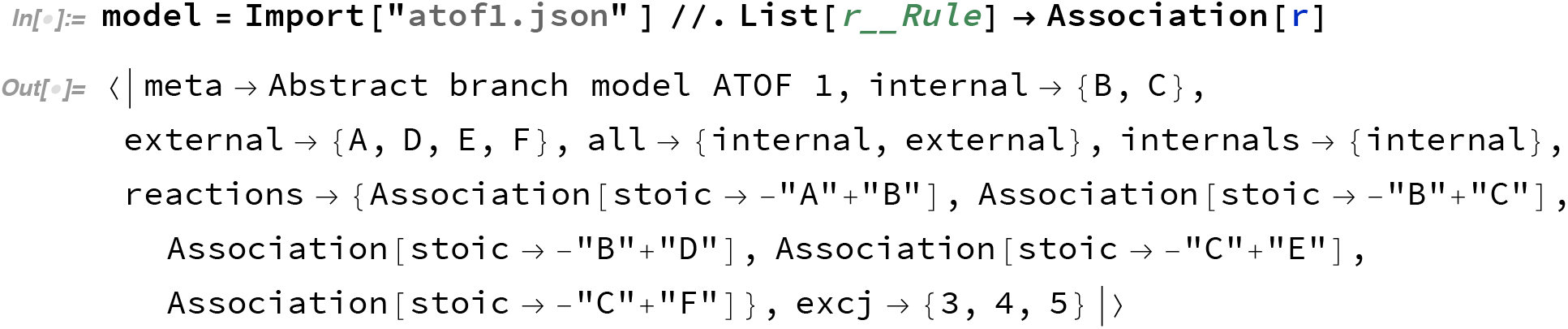

It has to be emphasized that the names of the metabolites: B, C, A, *etc*., are still character strings (instead of WL symbols) inside `model`. Presently, this is the way designed to keep the model definition file separated from the details of the computation.

In order to analyze the network structure, first the program needs to know the metabolites in the network. This is done by processing the `all` and `internals` fields in the JSON file and convert them into flat (1-dimensional) lists:

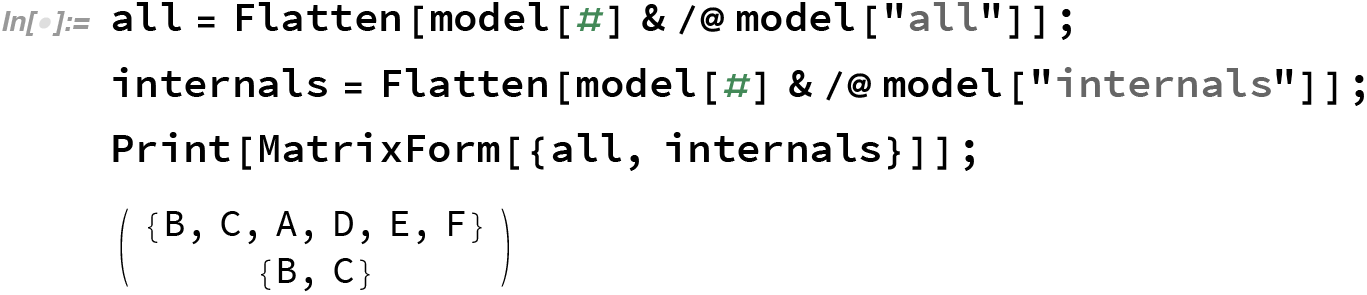

Notice that the order of the metabolites are strictly consistent with that given in the JSON file. This is usually a good thing, because it can be seen from the JSON file that metabolites will be imported in groups that the user defined. That means, metabolites with related properties can be arranged in the way the user desires to as much as possible. According to Stephanopoulos, for analyzing the independent pathways and network structure, only the internal metabolites and the steady-state internal metabolite stoichiometric coefficient matrix are necessary. However, for other analyses, it is often necessary to include all metabolites. In this article, for demonstration purpose, all of the metabolites are imported. Nevertheless, what does it mean by ‘all’ is determined by the model definition file.

In any case, the number of metabolites included will only affect the first dimension of the stoichiometric coefficient matrix. The second dimension of the stoichiometric coefficient matrix, which corresponds to the number of elementary fluxes considered in the network, remains the same in both cases. Notice that the independent pathways or the grouped pathways that shall be explored later all consist of the elementary fluxes given in the model. Therefore, even if in the present program not all of the metabolites were considered, after analysis in the following procedures, the results can all be carried over to the complete stoichiometric coefficient matrix for other analyses.

Next we need to construct the complete stoichiometric coefficient matrix (SCM) and the steady-state internal metabolite stoichiometric coefficient matrix (SIMS). To efficiently discuss the procedure, some special notations are introduced in the following text. To represent a column vector, say named *c*, here the notation *|c*) is used. This is a novel notation introduced by the author itself. However, most people familiar with quantum mechanics could possibly understand that the author was inspired by Dirac notation when introducing this new notation. Thus, it is not surprising to find that the author shall use the notation (*r|* to represent a row vector named *r*. Moreover, for a matrix, say named *A*, of any size, even when one of its dimensions is 1, as long as it is considered as a matrix, it will be represented as 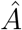. This notation is also parallel to the linear operators in many quantum mechanics literature.

Suppose *M* metabolites and *N* fluxes (bidirectional reactions) are in the model. The SCM, written as 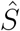, will be an *M × N* matrix. If the *M* metabolites are the set of all internal metabolites, this SCM is also the SIMS, written as 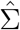. Therefore in the following we simply talk about SCM, and the discussions apply to SIMS. 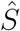 can also be considered as a row of *N* column vectors, each of dimension *M* and represents one flux:

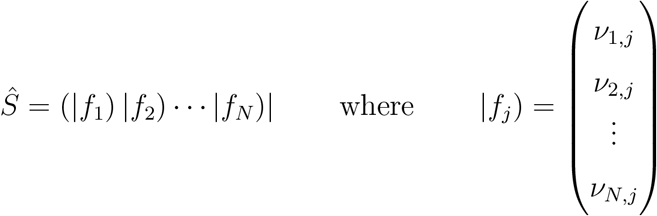

and *ν*_*i,j*_ is the stoichiometric coefficient of the *i*-th metabolite in the *j*-th flux. Therefore, to generate the correct SCM, we first need to put the right coefficient at the right place in a column for a given reaction. To that end, first import the description of all the reactions to see what is obtained:

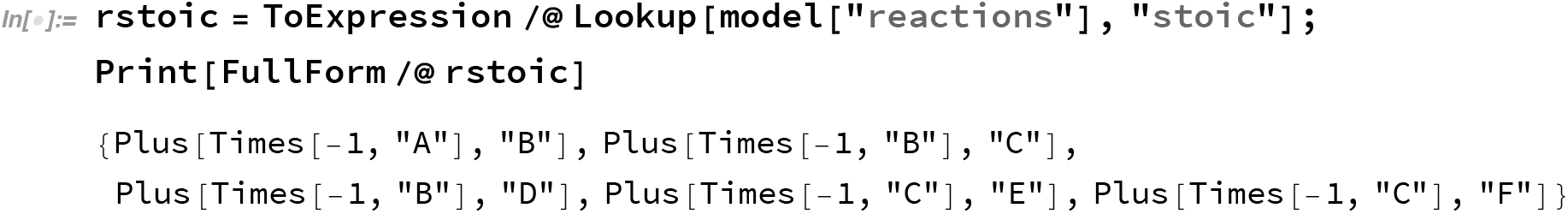

The full form of each item in the list is shown so that one can see that, even after converting the stoichiometric equation to WL expression, the names of the metabolites are still the original character strings. The trick used to convert one of the fluxes into a column is as following. First a column *|*`avec`) of dimension *M* is constructed, of which the *j*-th component is a `List` with only one `Rule` relating the name of the *j*-th metabolite to 1.

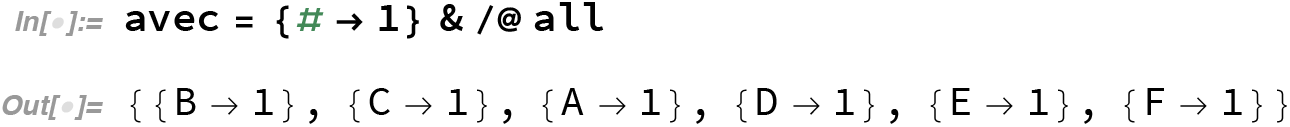

Remember that in WL a one-dimensional vector is always a column vector by default, although it is shown in a row to save space. It is easier to see the usage of `avec` than to say it:

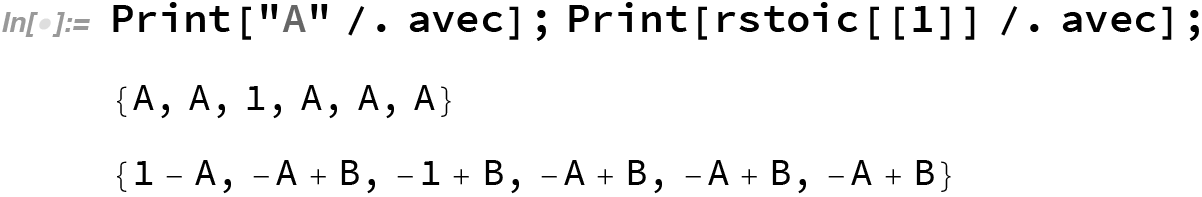

In the first example, character string `"A"` is **`ReplaceAll`** with avec. Since in avec the rule to convert string `"A"` to 1 is the 3rd component, after the replacement operation the result is a *M*-component vector where the 3rd component becomes 1, while all other components are still `"A"` because those components do not contain rule to convert string `"A"`. In the second example, the first component of `rstoic` (the stoichiometric equation of reaction 1, namely, `-"A"+"B"`) is **`ReplaceAll`** with `avec`. Since in `avec` only the first component contains rule to convert `B` to 1, the 1st component of the result becomes `1-"A"`. `"A"` is still there because there is not a rule to convert it in this component. Similarly, the 3rd component of the result becomes `-1+"B"`. Notice that since the coefficient of `"A"` is *−*1, replacing `"A"` by 1 results in a *−*1 in the place of `"A"`. All the other components besides the 1st and the 3rd are still the same as `rstoic[[1]]` because they cannot convert `"A"` or `"B"`. Thus, the (almost) correct vector is obtained after applying **`ReplaceAll`** to this stoichiometric equation with `avec`. As the final step, everything in the above result that remains as a character string is replaced with 0 and one can obtain the required result. One can directly apply the above operations on `rstoic`:

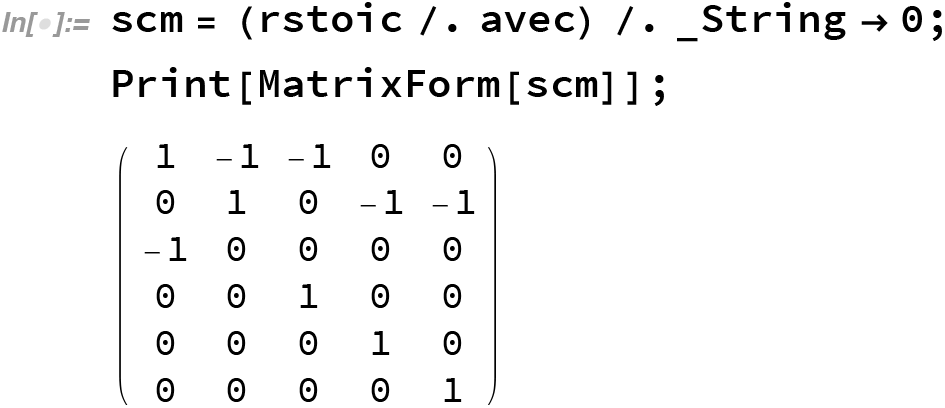

To obtain SIMS, one simply construct `ivec` similar to `avec` and apply the conversions:

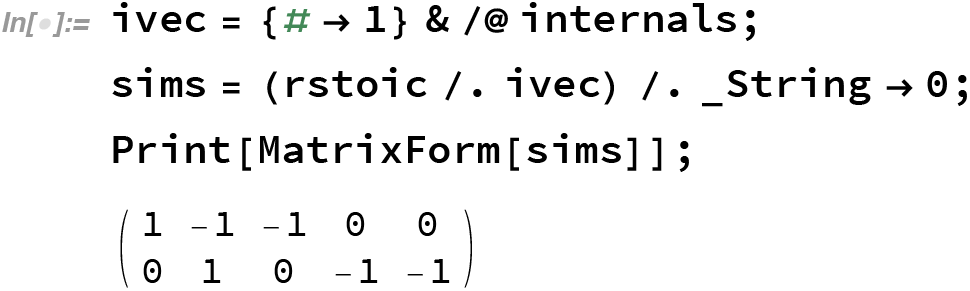

After obtaining the SIMS 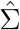, the independent pathways can be obtained by solving the equation homogeneous linear equation for the kernel 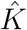:

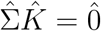

The dimension of 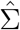 is *M × N* where *M* is the number of internal metabolites and *N* is the number of reactions. Therefore the first dimension (number of rows) of 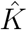 is also the number of reactions. The second dimension of 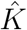 is the difference between the number of columns and the number of rows of 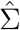. In their book, Stephanopoulos provided the reasoning in detail. Here it is only reminded that for a SIMS *M* is always less than *N*, but it is always full (row) rank. Therefore, there are always *N −M* free columns. Fundamental knowledge about linear algebra is sufficient to make sure that the null space of 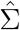 consists of *N − M* vectors that are *N*-dimensional.[5] Consequently, the zero matrix 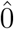 should be an *M ×* (*N − M*) matrix of 0.

However, the built-in function, **`NullSpace`**, of WL cannot be directly applied to obtain the 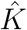 useful for further analysis. This is not because the null space provided by WL or other software like MATLAB is wrong. Null space is not unique in the first place. The kernel 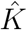 is also a representation of the null space. But there are two major constraints. First, the user may have an idea of which *N − M* of the exchange fluxes contribute only to one independent pathway each. Therefore, in a row corresponding to one of the correct exchange fluxes, all but one matrix elements are zero. The only non-zero component should be 1. If the correct exchange fluxes are chosen as the conditions to fix the free rows in the kernel, one correct kernel will be obtained, and it shall have the good property that all of the matrix elements of the kernel are non-negative. The second constraint on kernel 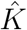 is exactly that all matrix elements of it are non-negative. Moreover, if the exchange fluxes were not assigned correctly, a kernel may not be found.

Take the model ATOF 1 as example, it is quite obvious that the fluxes 3, 4, and 5 are independent free fluxes. Therefore, first a 5*×*3 (or *M ×N*) matrix with all of the elements are unknown is constructed. Then, rows 3, 4, and 5, which form a 3*×*3 (or (*N − M*)*×*(*N − M*)) square matrix, are replaced by a 3*×*3 unit matrix. Thus, a matrix with 2 rows (or *M* rows) of unknown matrix elements, the row indexes of which are in the variable intjs, is constructed:

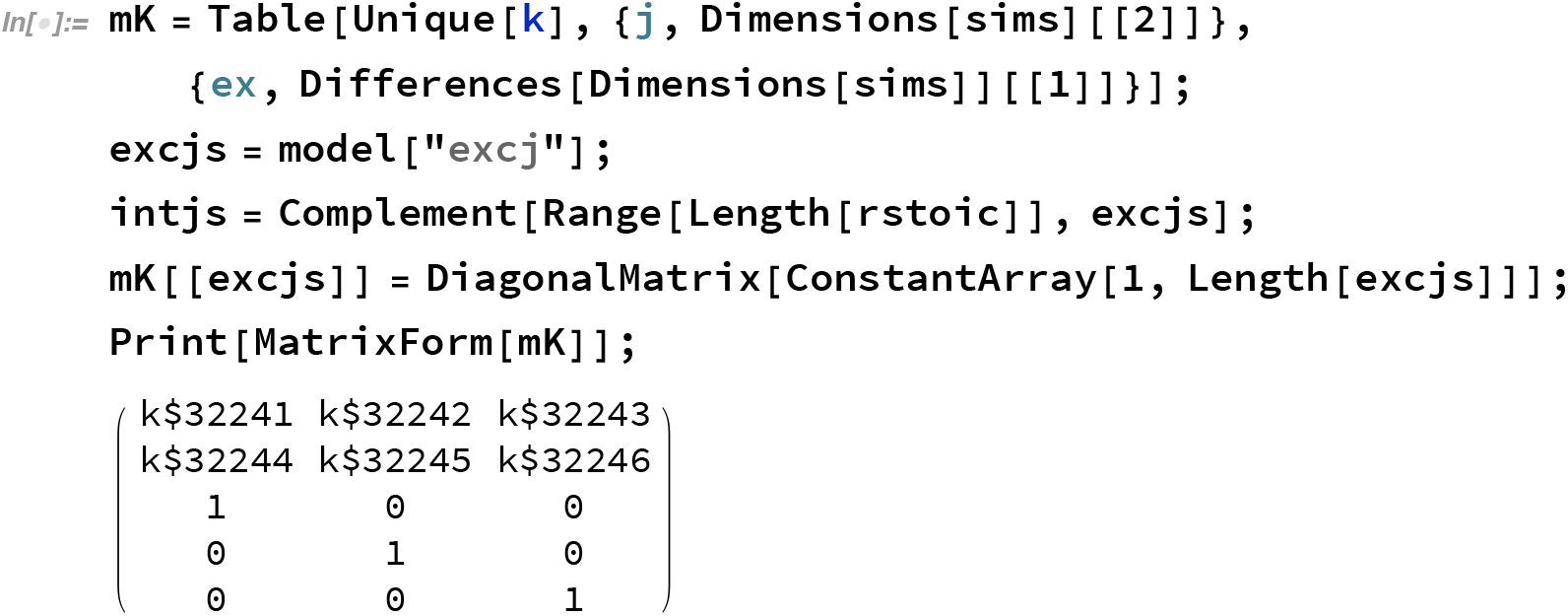

Therefore, the equation 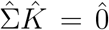 becomes a set of *M ×* (*N − M*) equations (the number comes from the dimensions of the zero matrix) with *M ×* (*N − M*) unknowns. Solving this set of equations, one can find the kernel:

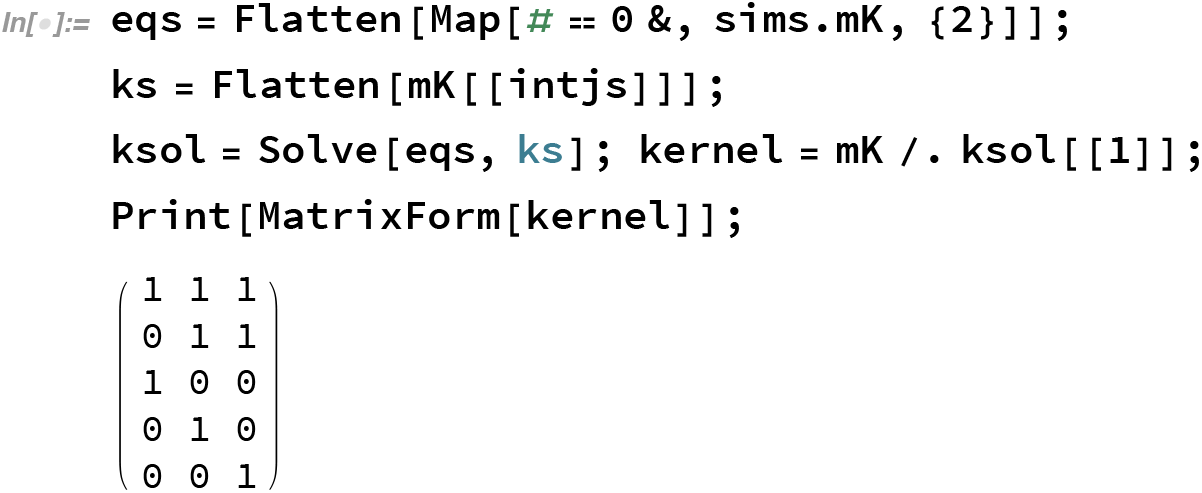

The three columns of 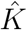 are the three independent pathways. They are shown as the dashed lines in the following diagram.

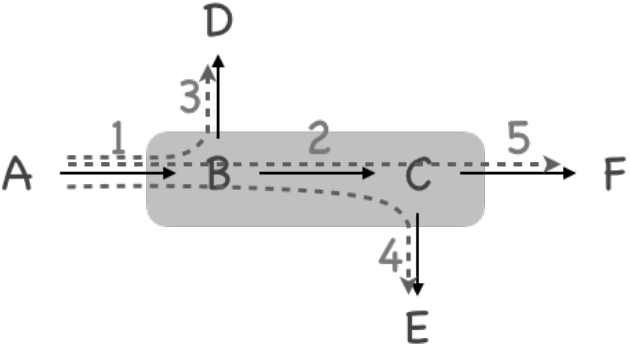

As summary, in this first step of structure analysis, the result of analyzing model ATOF model 1 is:

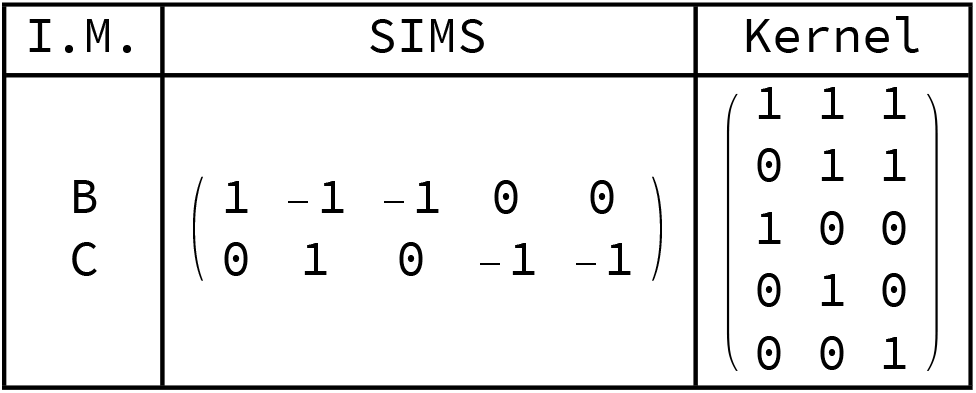
The pathway analysis of model ATOF 1.

Here I.M. is the abbreviation of internal metabolites.

There is not a problem applying the above procedures to other networks except that if one cannot determine the correct exchange fluxes the procedures would fail. Unfortunately, by far, there does not seem to be a clear mathematical algorithm to figure out the correct exchange fluxes to use without artificial intervention. It is hoped that such an algorithm can be developed in the near future. In this article we can only show some of the observations which may help the advance of the theory.

In the SIMS of ATOF 1, fluxes 3, 4, and 5 were finally decided to be the correct exchange fluxes to use. With hindsight, there is only one independent pathway going through each of them. However, automation of the procedures means exactly that hindsight or intuition cannot be used for any of the judgments in the flow chart. There must be a logic which can be converted to mathematical-logical operations. Looking at the SIMS of ATOF 1. It is found that all of the non-zero components of column 3, 4, and 5 are negative.

Let us apply the above program fragments to the aromatic amino acid synthesis pathway (AAASP) and check its SIMS:

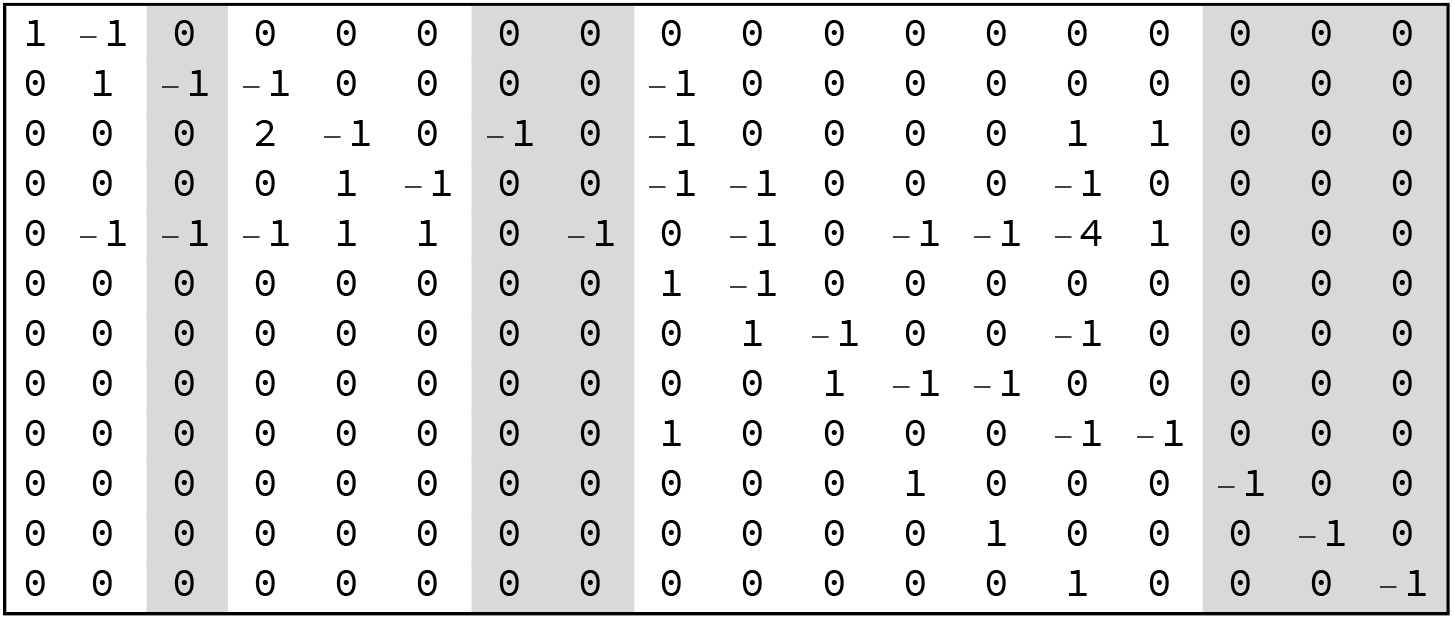
The SIMS of AAASP model.

It is found that only column 3, 7, 8, 16, 17, and 18 has such properties. It turns out that indeed they are the proper exchange pathways to choose. However, from the diagram of the model, it is found that seemingly fluxes 1, 3, 6, 7, 8, 16, 17, and 18 are all terminal fluxes. Why, then, are fluxes 1 and 6 not good exchange fluxes? For flux 1 it is obvious that the sign of its only nonzero component is positive, indicating that it only uptake the (major) substrate(s) and increases the content(s) of the substrate(s) in the system. In that case, it is a flux which most if not all independent pathways should go through. It may not be a good choice as exchange flux for this reason. For flux 6, it seems (misleadingly) that it only consumes PEP. However, the plus sign next to it indicates that it also produces ATP. In that case, through flux 8, flux 6 will be coupled back to other reactions in this network. It is actually not a terminal flux. Therefore it seems quite inspiring to us that this rule of non-positive column may be used to pick the correct exchange fluxes.

Unfortunately, if one looked at the SIMS of the ATOF 2 model, this rule does not hold. Let us look at the result of complete independent pathway analysis of ATOF 2:

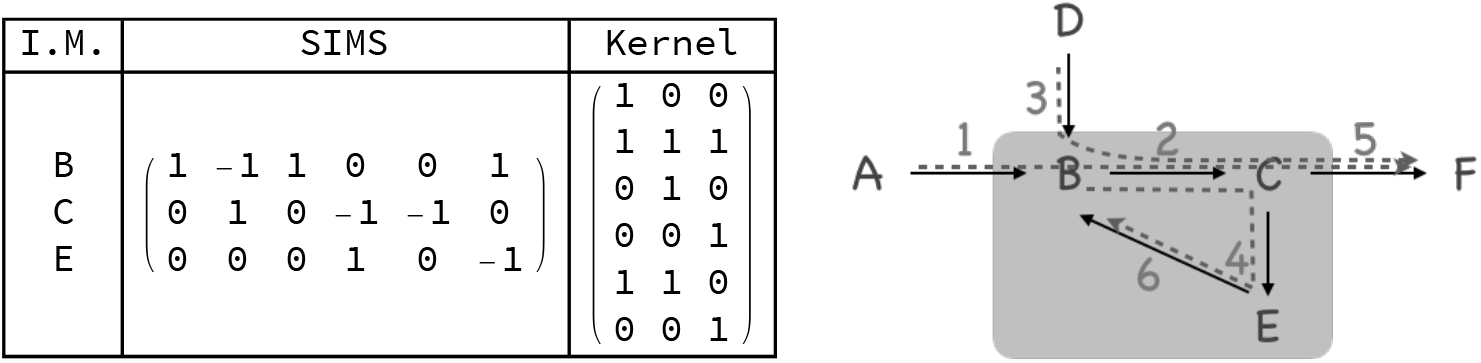
Independent pathways analysis of ATOF 2 model.

For ATOF 2, fluxes 1, 3, and 4 were chosen as exchange fluxes. This can be guessed from the final expression of the kernel. However, the analysis results is exactly the same (although the details of all steps changed slightly) if one chose fluxes 1, 3, and 6 instead. With hindsight, it can be seen that fluxes 4 and 6 are indeed equivalent, and they are both traversed only by one and the same independent pathway. The trouble is, they are neither terminal flux. They link the internal metabolites back to the network.

One may be tempted to say that problems would arise when there are some “totally internal independent pathways”. Then the question is: how could one detect such totally internal independent pathways just by looking at the SIMS? And, if internal independent pathways were located, what is the algorithm to pick a proper exchange flux from there? This is an unsolved problem of the present work. Nevertheless, to be optimistic, in realistic networks, similar to AAASP, each flux may often involve more than one input or output internal metabolites. This is called the nonlinearity of the flux.[3] In realistic networks, totally internal independent pathways may be less likely to appear. It is one’s best hope, for now. One can try to pick out columns with completely non-positive components and check if there are *N − M* of them. If there are not enough such columns, the procedure can go on without problem. If not enough such columns (fluxes) were picked, one has to investigate the network partially manually.

## IV. LINK METABOLITE ANALYSIS

The next goal of metabolic networks structure analysis is to locate the branching points in the networks where independent pathways diverse from each other. The internal metabolites on these network-topological branch points are called the *link metabolites* (LM).

Since branching is only meaningful for those independent pathways found in the analysis in the previous section, the following procedures start with and mainly focus on the kernel matrix 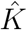. Of course some properties of SIMS of course are crucial for the analysis. Here, an information very useful for the LM analysis is the internal metabolite(s) involved in each flux. In this work, the following method is used to obtain this information.

First, all of the nonzero elements of matrix 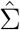 are converted to 1. Second, construct an *M × M* diagonal matrix with the list of internal metabolites (in this program they are character strings of names) as the diagonal. Multiply the name diagonal matrix to the modified SIMS from the left, an *M × N* matrix showing which IM appears in which flux is obtained. For convenience, transpose this matrix so that there are *N* rows and *M* column in it. For each row, which is a vector, but can also be considered as a set, remove 0 from it. Then the above matrix becomes a column of *N* sets, each corresponds to one flux, and contains the IMs involved in this flux. This variable is called `inms`. Take ATOF 1 as the example, the operation is as following:

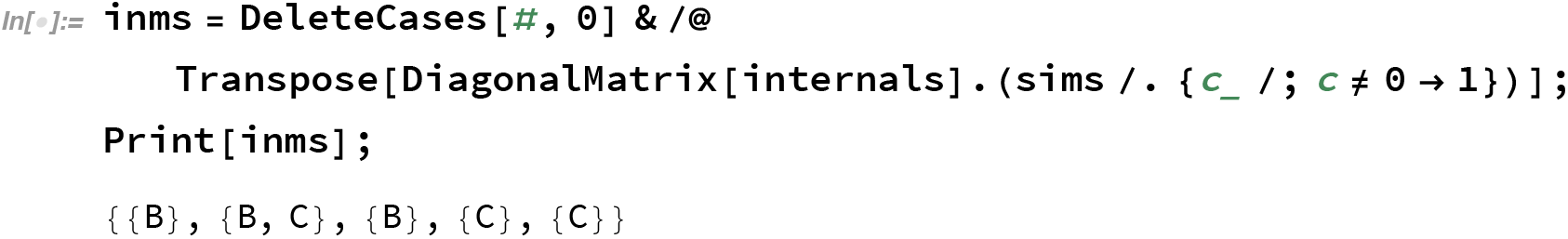

Similarly, the `inms` for ATOF 2 and AAASP can be obtained as following:

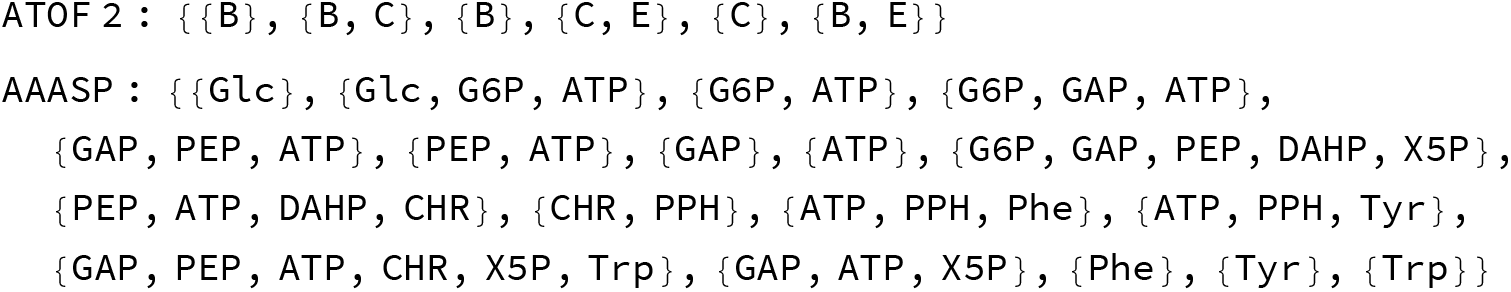

This information will be used later.

According to Stephanopoulos, the first step to analyze the kernel is to find the flux(es) common to all of the independent pathways. Mathematically, a common flux is a row in 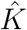 whose components are all nonzero. To test whether a row contains zero or not, simply multiply all its components together. The product will be zero unless it is a common flux. In the ATOF 1 model:

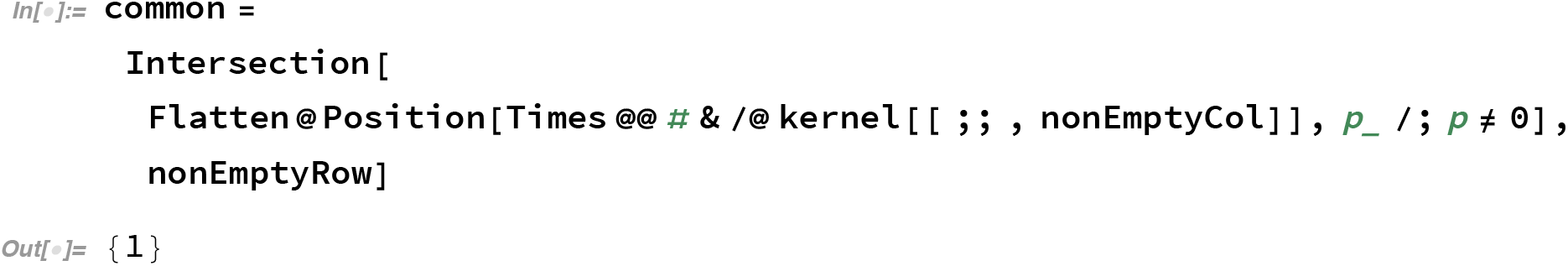

Notice that there are two variables not explained by far. The `nonEmptyCol` contains the indexes of all of the columns that are not all 0; Similarly `nonEmptyRow` contains the indexes of all of the rows that are not all 0. At this stage, they contain all the columns and all the rows, respectively. How they are changed during the analysis will be explained later. As the result, the first flux is picked out. Looking at the kernel matrix of ATOF 1 the result can be directly understood:

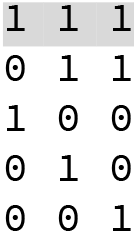
The kernel matrix of ATOF 1.

The product of flux 1 is B, which is the only internal metabolite involved in flux 1. Therefore there is no difficulty to select the link metabolite. It must be B. In complicated networks like AAASP, the choice of LM is much more complicated. But here the procedure will be outlined with the simple example first. The complications will be discussed afterwards.

The common fluxes just found is called the group A pathway in Stephanopoulos’ book. At the branch point, the fluxes will be separated into two. One is usually a terminal independent pathway, and is called the group B pathway; The other one is usually a set of several other independent pathways, and is called the group C pathway. The magnitude (coefficients) of group A should be determined, and then group A and group B will not be further analyzed. group C will become the next ‘kernel’ to be further analyzed.

Now let us at the `inms` of ATOF 1. The first flux is already identified as the group A pathway so it is ignored at the moment. In other fluxes, only flux 3 involves and only involves metabolite B. Looking at the kernel matrix except for the common rows (which is only row 1 at the moment), only the first column has a contribution from flux 3. Therefore, the first independent pathway is identified as the group B pathway.

The group B pathway should be eliminated in order to obtain the kernel for finding the next LM. The goal of elimination is to make all other components in the last (production) row of the common fluxes. After the elimination, a new matrix is obtained, and it is the group C pathway as well as the kernel for the next round of analysis, to find the next branch point and link metabolite. In Stephanopoulos’ book, these steps are described in great detail. Therefore it will not be further elaborated here. The steps for this round of analysis is summarized in a diagram for reference.

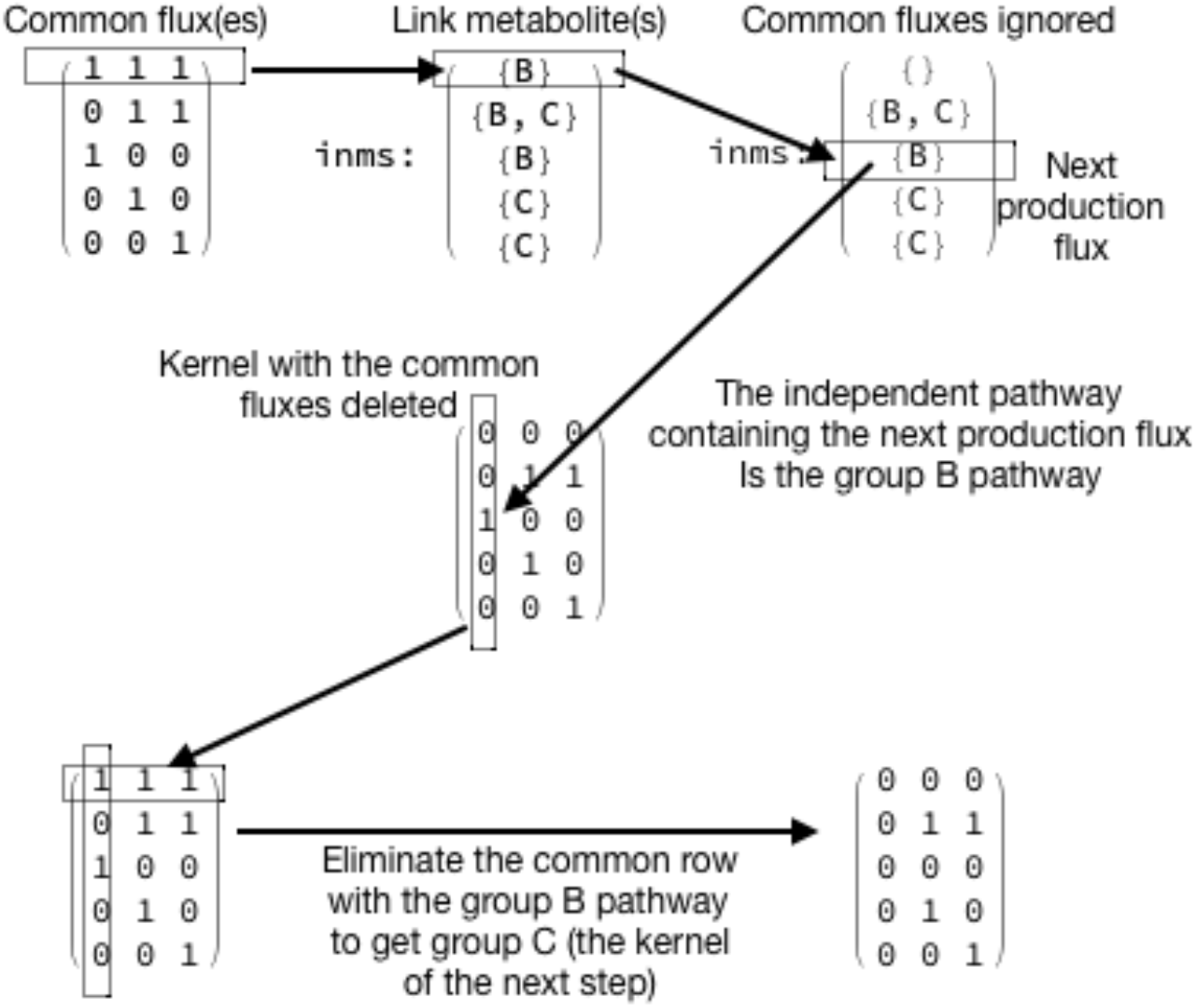

In ATOF 1 model, there are three independent pathways. Therefore, two branch points are necessary to completely separate the three pathways. Therefore, in general, the number of branch points is *N − M −* 1. That means, the above analysis has to be repeated *N − M −* 1 times. When going into the next round, except finding the new kernel matrix for the analysis, which has just been done, one also need to eliminate the common independent pathways that were already separated into group B of each step. The link metabolites identified in previous rounds also have to be discard in later rounds: one metabolite cannot be the link metabolite in two branch points. Also, during the analysis, it has to be careful not to work on rows that are already totally eliminated. Avoiding those all-zero rows will make the program much safer to use. These are details that have to be covered by the program. Here the implementations of these details are not described in further detail unless absolutely necessary. The loop code that handles all of the analysis steps is shown below.

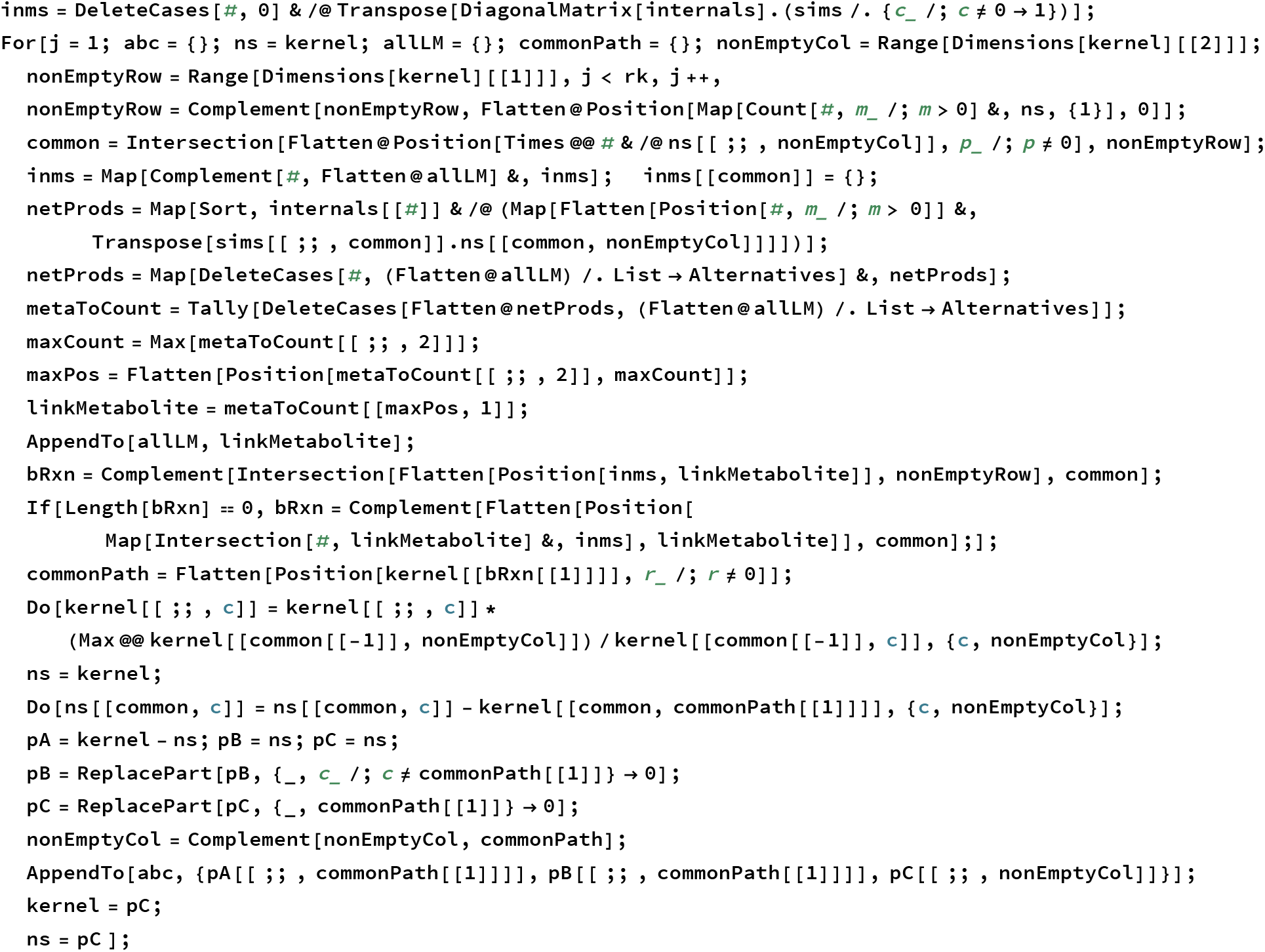

The results, including the link metabolite and the group A, group B, and group C matrices obtained in each step can be shown. The index of the round is shown in the first row in the table of each model. The name of the internal metabolite in each round is in the second row. The group A, group B, and group C pathways are shown as matrices in due order in the third row. Below are the results for analyzing ATOF 1 and 2, respectively:

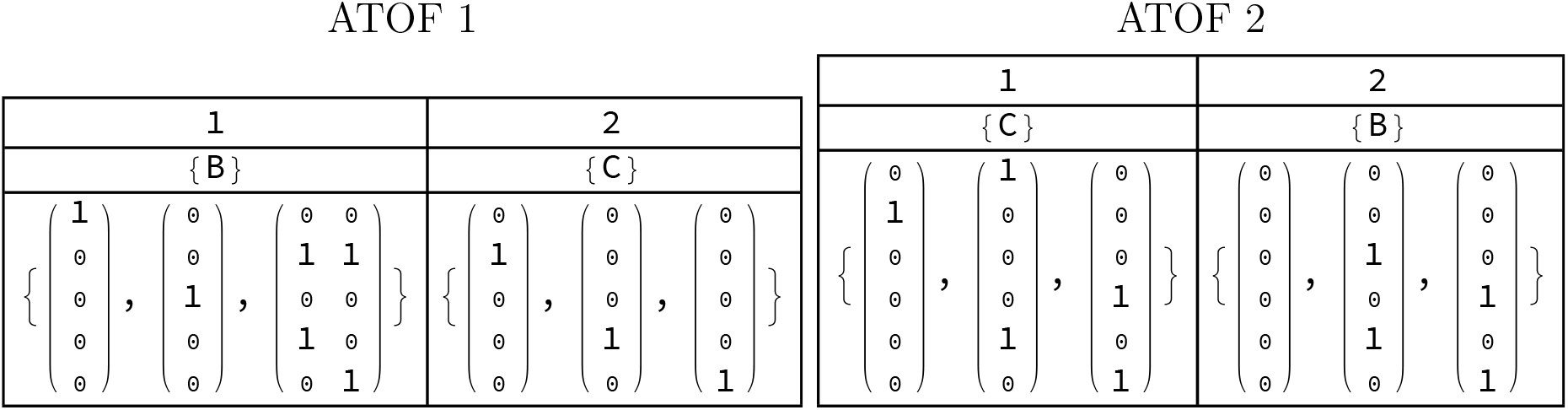

The remaining problems are: for complicated networks like AAASP, there are steps in which the simple approach described for ATOF 1 should actually be somewhat more complicated. In the following, the analysis of the AAASP model is shown, but not all the details will be repeated. Only the occasion at which the AAASP case is different from the ATOF 1 case is discussed. That is supposed to help readers understand why the WL code is more complicated than described in the ATOF 1 case.

Let us start with the fundamental properties of the AAASP model.

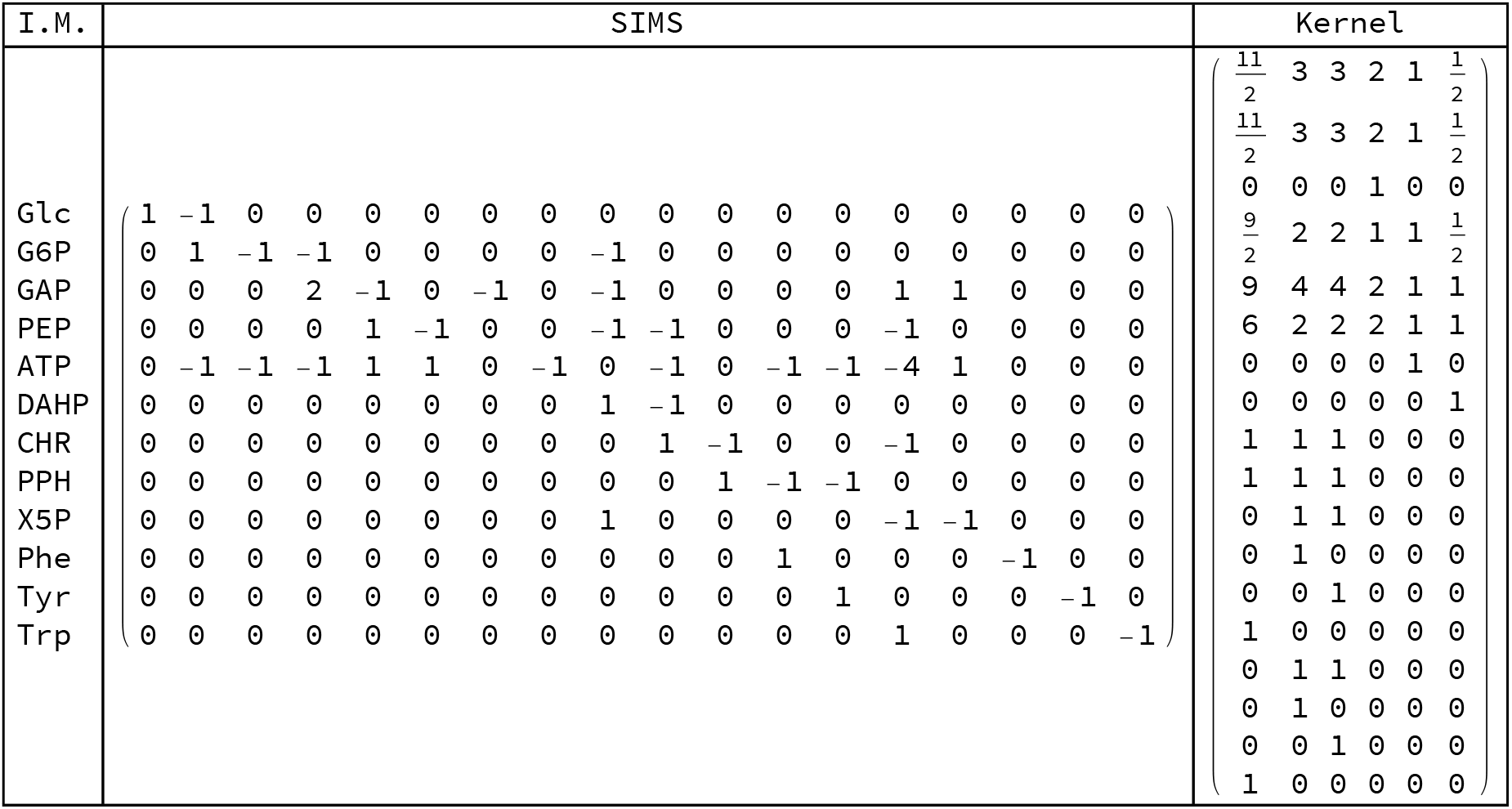

The `inms` is already shown earlier. It is found that the fluxes 1, 2, 4, 5, and 6 are common fluxes in the first round. The last flux, that is, flux 6, has a net production of only ATP. This is a strong hint that ATP is the link metabolite. However, the situation may be much more complicated than that. If we continued to pick the net product of only the last common reaction, it works for the first three round. ATP, G6P, and GAP will be identified as the LM for the first three round. But upon proceeding to the 4th round of analysis, the common fluxes will be fluxes 5, 9, and 10. Flux 10 produce CHR, and flux 9 has net production of X5P and DAHP. In the diagram of AAASP, fluxes 5, 9, and 10 form very complicated cycle. Actually, from the diagram, one can hardly find a reason to explain why flux 9 has to be arranged before flux 10. Their order can be freely reversed. In contrast, the order of fluxes 1, 2, 4, 5, and 6 can hardly be changed. If changed, there will be problem since the early round. For example, if flux 6 is not the last flux in the common fluxes in round 1, there will be trouble finding the first LM.

Moreover, how ever one rearrange the order of fluxes 9 and 10, problem remains. If one selected CHR as the LM, he would obtain two independent pathways as group B. Similarly if one selected X5P and DAHP as LMs, the analysis fails to proceed.

Actually, there is fundamental reasons behind the problem. Choosing the net product of the last common flux is by itself not the correct strategy.

Earlier, it is explained that the kernel matrix is the null space of SIMS. However, if only part of the *N* reactions are considered, that is, if one took several columns from SIMS, and take the rows of the kernel with corresponding indexes, and multiply them together, the result is still an *M ×* (*N − M*) matrix, but this time it will not be totally zero. This simply reflects the trivial fact that the null space of the whole network is not the null space of part of the network. Thus, if a part of the network, that is, the common fluxes, were picked, one can find the overall net products of the subnetwork. This is done by the following operation:

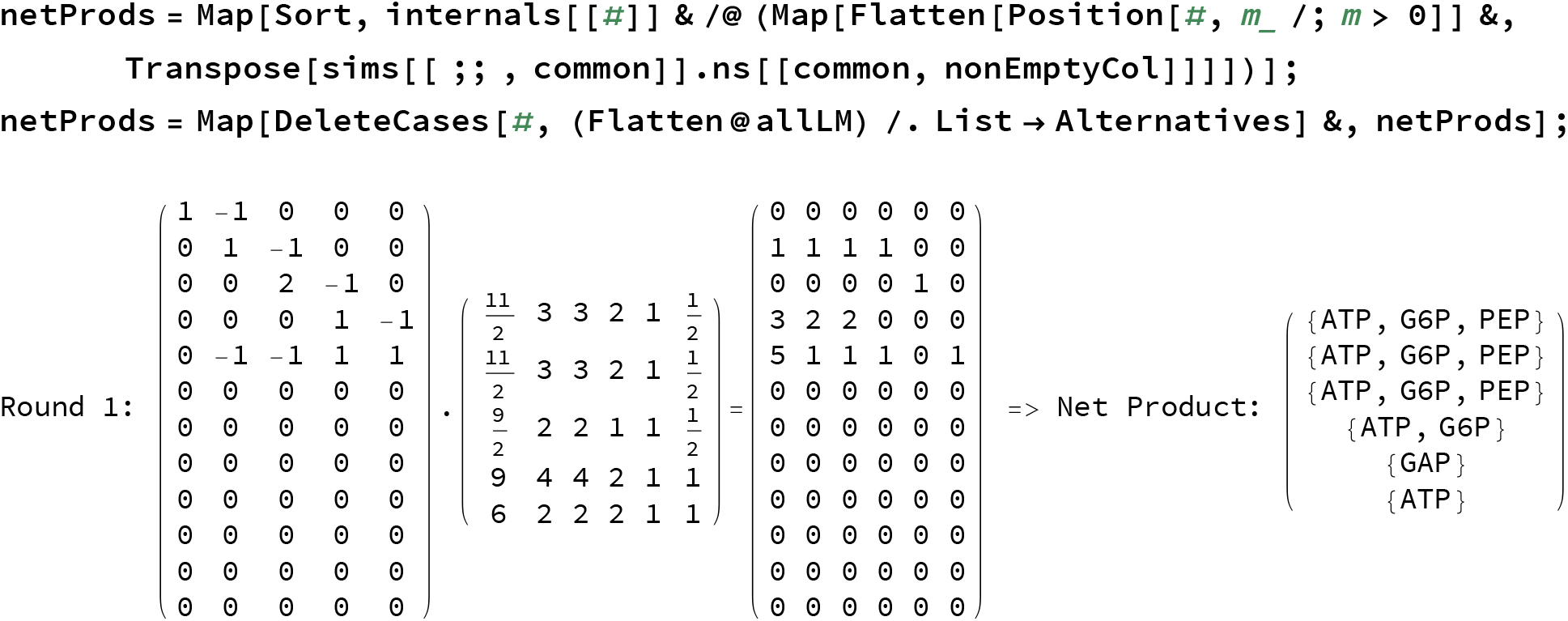

In the resulted variable `netProds`, each component represents the net products of the subnetwork, if the weights on each flux (the common fluxes) is in accordance with those in the corresponding independent pathway in the kernel. For example, in the 6th column of the product of the two partial matrices, there is only a positive component at the 5th row. That means, following the weight of the 6th independent pathway, this subnet only has net production of ATP, which is the 5th metabolite. However, one can not say that ATP is the link metabolite yet. Of the six independent pathways, one of them does not produce ATP. Independent pathway number 5 produces only GAP. Honestly speaking, the author does not know for sure what would happen if GAP is chosen as the link metabolite. The only thing that is sure is that the result will be quite different from that given by Stephanopoulos.

Instead of choosing the net product from one of the pathways, which does not seem to be a reasonable approach (because the order of the independent pathways is also arbitrary), here the link metabolites is decided as the metabolite(s) that appear(s) in the most number of independent pathways that has to be considered. It is found that ATP appears in 5 of the 6 independent pathways, therefore, it is the choice in this round. For information, in the second round G6P is the only candidate of LM; In the third round, GAP is the only candidate of LM; In the fourth round, CHR and X5P both appear in all three remaining independent pathways while PEP appears once, and the appearance of DHAP happens to cancel out in all three pathways so it never appears; In the 5th and the final round, only PPH appears in the two remaining independent pathways.

The determination of link metabolite(s) is perhaps the most difficult part of the task. Afterwards, when one tries to select the flux that belongs to group B, there is also complication. In the `inms` variable, there may not be any component that exactly match the link metabolite. In that case the program cannot determine the group B pathway. What is done in the program is that the component(s) of `inms` that include all of the link metabolites are picked. In this compromised way, often more than one components are picked. But according to experience, in such cases the choice can then be arbitrary. The program simply picks the first component satisfying the criteria.

Finally, the result of the five round of analysis of the AAASP model is shown below.

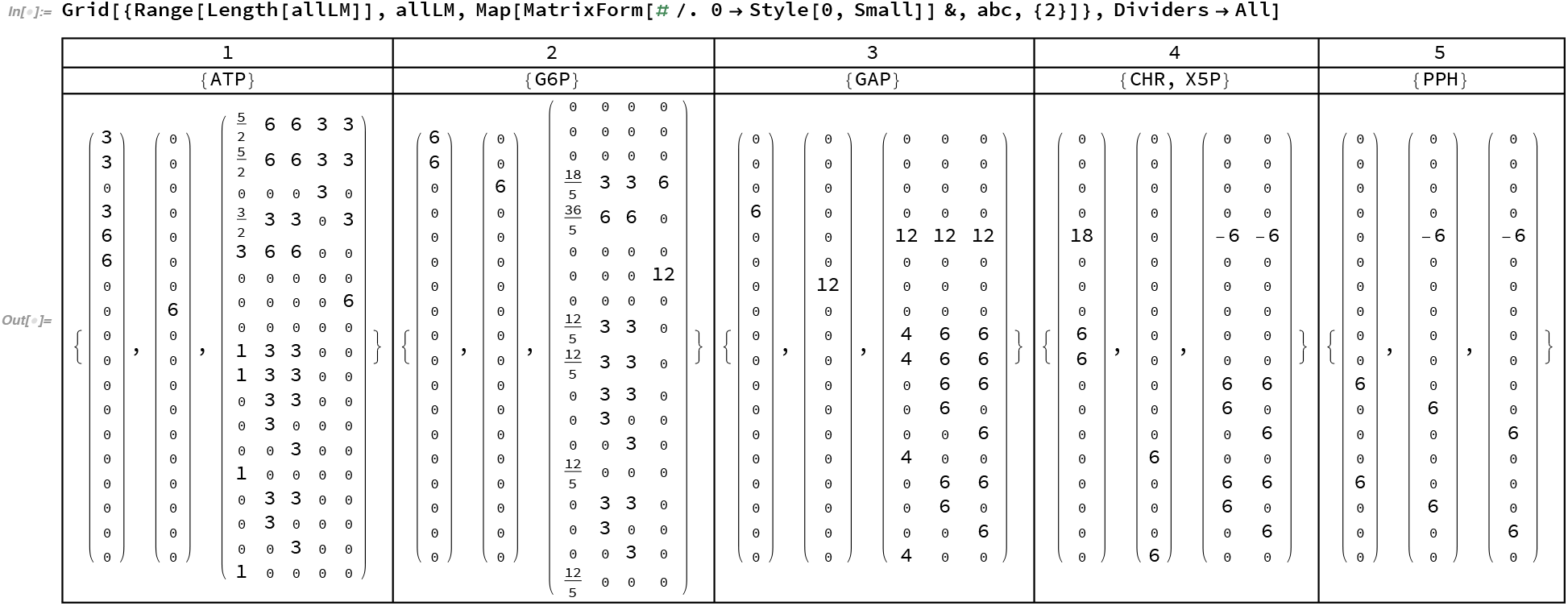

## V. CONCLUSION

In this work, a totally automatic program that can analyze the structure of a metabolic network is presented. This problem of network structure analysis is actually quite complicated to be automated compared to many other further analyses (metabolic flux and control analysis, and so no.). It is difficult to explain the details of a program by words. The author welcomes further discussions and cooperation to explore other automatic analyses.

